# Optogenetic stimulation of glutamatergic neurons in the cuneiform nucleus controls locomotor movements in a mouse model of Parkinson’s disease

**DOI:** 10.1101/2021.06.13.448213

**Authors:** Maxime Fougère, Cornelis Immanuel van der Zouwen, Joël Boutin, Kloé Neszvecsko, Philippe Sarret, Dimitri Ryczko

## Abstract

In Parkinson’s disease (**PD**), the loss of midbrain dopaminergic cells results in severe locomotor deficits such a gait freezing and akinesia. Growing evidence indicates that these deficits can be attributed to decreased activity in the Mesencephalic Locomotor Region (**MLR**), a brainstem region controlling locomotion. Clinicians are exploring deep brain stimulation of the MLR as a treatment option to improve locomotor function. The results are variable, from modest to promising. However, within the MLR, clinicians have targeted the pedunculopontine nucleus exclusively, while leaving the cuneiform nucleus unexplored. To our knowledge, the effects of cuneiform nucleus stimulation have never been determined in parkinsonian conditions in any animal model. Here, we addressed this issue in a mouse model of Parkinson’s disease based on bilateral striatal injection of 6-hydroxydopamine (**6-OHDA**), which damaged the nigrostriatal pathway and decreased locomotor activity. We show that selective optogenetic stimulation of glutamatergic neurons in the cuneiform nucleus in mice expressing channelrhodopsin in a Cre-dependent manner in Vglut2-positive neurons (**Vglut2-ChR2-EYFP mice**) increased the number of locomotor initiations, increased the time spent in locomotion, and controlled locomotor speed. Using deep learning-based movement analysis, we found that limb kinematics of optogenetic-evoked locomotion in pathological conditions were largely similar to those recorded in freely moving animals. Our work identifies the glutamatergic neurons of the cuneiform nucleus as a potentially clinically relevant target to improve locomotor activity in parkinsonian conditions. Our study should open new avenues to develop targeted stimulation of these neurons using deep brain stimulation, pharmacotherapy or optogenetics.

**SIGNIFICANCE STATEMENT:** In Parkinson’s disease, alleviating locomotor deficits is a challenge. Clinicians are exploring deep brain stimulation of the Mesencephalic Locomotor Region, a brainstem region controlling locomotion, but results are mixed. However, the best target in this region in Parkinson’s disease remains unknown. Indeed, this region which comprises the pedunculopontine and cuneiform nuclei, contains different cell types with opposing effects on locomotor output. Here, using a mouse model where midbrain dopaminergic cells were damaged by a neurotoxin, we demonstrate that optogenetic activation of glutamatergic neurons in the cuneiform nucleus increases locomotion, controls speed, and evokes limb movements similar to those observed during spontaneous locomotion in intact animals. Our study identifies a potentially clinically relevant target to improve locomotor function in Parkinson’s disease.

## INTRODUCTION

In Parkinson’s disease (**PD**), midbrain dopaminergic (**DA**) cells are lost, resulting in motor dysfunction including severe locomotor deficits (e.g. gait freezing, akinesia, falls) (Bloem et al. 2004). Growing evidence indicates that part of these deficits can be attributed to changes in the Mesencephalic Locomotor Region (**MLR**) (Hirsch et al. 1987, Zweig et al. 1987, Jellinger 1988, Rinne et al. 2008, Demain et al. 2014, Karachi et al. 2010, for review Ryczko and Dubuc 2017). This brainstem region plays a key role in locomotor control by sending projections to reticulospinal neurons that carry the locomotor drive to the spinal cord in vertebrates (lamprey: Sirota et al. 2000, Brocard et al. 2010; salamander: Cabelguen et al. 2003, Ryczko et al. 2016a, mice: Bretzner and Brownstone 2013, Lee et al. 2014, Roseberry et al. 2016, Capelli et al. 2017, Caggiano et al. 2018, Josset et al. 2018, van der Zouwen et al. 2021; for review Ryczko and Dubuc 2013). The DA neurons of the *substantia nigra pars compacta* (**SNc**) indirectly control MLR activity through the basal ganglia (Stephenson-Jones et al. 2011, Kravitz et al. 2010, Roseberry et al. 2016). In parallel, the MLR receives direct DA projections from the SNc (Ryczko et al. 2013, 2016b, 2017, Perez-Fernandez et al. 2014) and from the zona incerta (Sharma et al. 2018). The DA innervation of the MLR degenerates in a monkey model of PD (Rolland et al. 2009). Therefore, the loss of DA cells in PD has major effects on MLR activity. In PD, locomotor deficits are associated with MLR cell loss, abnormal neural activity, altered connectivity, and metabolic deficits, likely resulting in a loss of amplification of the locomotor commands (for review Ryczko and Dubuc 2017). Accordingly, motor arrests and gait freezing are associated with a decrease in MLR activity in PD (Shine et al. 2013, for review Ryczko and Dubuc 2017).

One approach to improve locomotor function in PD would be to increase MLR activity. L-DOPA, the gold standard drug used to improve motor symptoms in PD, increases MLR activity and this likely contributes to the locomotor benefits (Fraix et al. 2013). The MLR has been proposed to contribute to the locomotor benefits of Deep Brain Stimulation (**DBS**) of the subthalamic nucleus (Holiga et al. 2015, Knight et al. 2015, Weiss et al. 2015), which has direct and indirect projections to the MLR (Breit et al. 2001, Neagu et al. 2013, for review Ryczko and Dubuc 2013). However, the benefits of L-DOPA and subthalamic DBS on locomotor deficits may wane over time, highlighting the need to find new therapeutic approaches (Sébille et al. 2019, Gavriliuc et al. 2020). Since 2005, the MLR is explored as a DBS target (Plaha and Gill 2005). The results vary, from modest to promising (Hamani et al. 2016a,b). However, the best target in the MLR in PD conditions is not yet identified. The MLR is a heterogeneous structure, with the cuneiform nucleus (**CnF**) controlling the largest range of locomotor speeds, and the pedunculopontine nucleus (**PPN**) rather controlling slow speeds, posture and in some cases locomotor arrests (Roseberry et al. 2016, Caggiano et al. 2018, Josset et al. 2018, van der Zouwen et al. 2021, for review Ryczko and Dubuc 2013). Human DBS protocols targeted the PPN, but left the CnF unexplored, despite its major importance in locomotor control in animal research (Chang et al. 2020, 2021b, van der Zouwen et al. 2021).

To add further complexity, three main cell types are present in the MLR: glutamatergic, GABAergic and cholinergic cells. It is still unknown which cell type is the best target to improve locomotor function in PD conditions. Optogenetic studies uncovered that glutamatergic cells in the CnF play a key role in generating the locomotor drive for a wide range of speeds (Roseberry et al. 2016, Caggiano et al. 2018, Josset et al. 2018, van der Zouwen et al. 2021). Glutamatergic cells of the PPN control slower speeds (Caggiano et al. 2018, Josset et al. 2018), and in some cases evoke locomotor arrests (Josset et al. 2018, Dautan et al. 2020). The GABAergic cells in the CnF and PPN stop locomotion likely by inhibiting glutamatergic cells (Roseberry et al. 2016, Caggiano et al. 2018). The role of the PPN cholinergic cells is not resolved, as their activation can increase or decrease locomotion (Roseberry et al. 2016, Caggiano et al. 2018, Josset et al. 2018 for review Ryczko and Dubuc 2013). Clinically, DBS likely stimulates all cells around the electrode, including the GABAergic cells that stop locomotion, and this could contribute to the variability of outcomes.

Here, we aimed at identifying a relevant target in the MLR to improve locomotor function in parkinsonian conditions. We hypothesized that selective activation of CnF glutamatergic neurons should improve locomotor function in a mouse model of PD. We induced parkinsonian conditions in mice by bilaterally injecting in the striatum the neurotoxin 6-hydroxydopamine (6-OHDA), which is well known to damage the nigrostriatal DA pathway and to induce a dramatic decrease in locomotor activity (e.g. Kravitz et al. 2010, Watson et al. 2021). Using *in vivo* optogenetics in mice expressing channelrhodopsin in a Cre-dependent manner in Vglut2-positive neurons (**Vglut2-ChR2-EYFP** mice), we show that photostimulation of glutamatergic neurons in the CnF robustly initiated locomotion, reduced immobility, increased the time spent in locomotion, and precisely controlled locomotor speed. Our results should help defining therapeutic strategies aimed at specifically activating CnF glutamatergic neurons to improve locomotor function in PD using optimized DBS protocols, pharmacotherapy or future optogenetic tools for human use.

## MATERIALS AND METHODS

### Ethics statement

All procedures were in accordance with the guidelines of the Canadian Council on Animal Care and were approved by the animal care and use committees of the Université de Sherbrooke (QC, Canada). Care was taken to minimize the number of animals used and their suffering.

### Animals

We used Vglut2-ires-Cre knock-in mice (Jackson laboratories, #028863, C57BL/6J) (Vong et al. 2011), ChR2-EYFP-lox mice (Ai32 mice, Jackson laboratory, #024109, B6.Cg-*Gt(ROSA)26Sor^tm32(CAG-COP4*H134R/EYFP)Hze^*/J) (Madisen et al. 2012), and ZsGreen-lox mice (Ai6 mice, Jackson laboratory, #007906, B6.Cg-*Gt(ROSA)26Sor^tm6(CAG-ZsGreen1)Hze^*/J) (Madisen et al. 2012). Homozygous Vglut2-ires-Cre knock-in mice were crossed with homozygous ChR2-EYFP-lox mice to obtain double heterozygous Vglut2-ChR2 mice. Homozygous Vglut2-ires-Cre knock-in mice were crossed with homozygous ZsGreen-lox mice to obtain double heterozygous Vglut2-ZsGreen-lox mice. Mice were genotyped as previously described (Fougère et al. 2021, van der Zouwen et al. 2021). Animals had *ad libitum* access to food and water, with lights on from 6 AM to 8 PM. Vglut2-ChR2-EYFP mice used for *in vivo* experiments were 22-46 weeks old at time of use (5 males, 6 females). Vglut2-ZsGreen mice used for RNAscope experiments were 5-8 weeks old (4 females), and those used for immunofluorescence control experiments were 6-18 weeks old (5 males, 1 female).

### PD model

The neurotoxin 6-OHDA was successfully used in mice to ablate DA neurons and evoke locomotor deficits resembling those reported in PD, including decreased number of locomotor initiations (Kravitz et al. 2010), increased freezing frequency (Kravitz et al. 2010) or akinesia (Watson et al. 2021). The mice received 30 min prior to 6-OHDA (or vehicle) injection, a systemic injection of desipramine to protect the noradrenergic and serotoninergic systems (25 mg/kg, 300 µL i.p.) (Thiele et al. 2012). Then, mice were anaesthetized using isoflurane (induction: 5%, 500 mL/min; maintenance: 1.5–2.5%, 100 mL/min) delivered with a SomnoSuite (Kent Scientific, Torrington, CT, USA) and placed in a Robot Stereotaxic instrument coupled with StereoDrive software (Neurostar, Tübingen, Germany). Two holes were drilled in the cranium and a 10 µL Hamilton syringe locked into the Robot Stereotaxic instrument was used to inject bilaterally in the striatum either 6-OHDA (0.2% ascorbic acid, 0.9% NaCl saline solution, 6-OHDA 5 mg/mL, 1 μL/side) or the vehicle (0.2% ascorbic acid, 0.9% NaCl saline solution, 1 μL/side) (Kravitz et al. 2010, Thiele et al. 2012). The syringe needle was lowered at +0.50 mm anteroposterior, +/-1.50 mm mediolateral, -3.00 mm dorsoventral relative to bregma. The compound injected was delivered at a rate of 0.2 µL/min. The syringe was left in place for 5 min before being removed. The scalp incision was sutured, and antibiotics were applied locally. The animals were tested 3 days after 6-OHDA injection, when the DA cells have degenerated, and locomotor deficits are observed (Kravitz et al. 2010, Watson et al. 2021).

### Optic fiber implantation

The procedure was as previously reported (van der Zouwen et al. 2021). Briefly, mice were anaesthetized using isoflurane (induction: 5%, 500 mL/min; maintenance: 1.5–2.5%, 100 mL/min) delivered with a SomnoSuite (Kent Scientific, Torrington, CT, USA). Mice were placed in a Robot Stereotaxic instrument coupled with StereoDrive software (Neurostar, Tübingen, Germany). An incision was made on the scalp, a hole was drilled in the cranium, and a fiber (200 mm core, 0.22 NA, Thorlabs, Newton, NJ, USA) held in a 5 mm ceramic or stainless-steel ferrule was placed 500 µm above the right CnF at -4.70 to 4.75 mm anteroposterior, +1.15 mm mediolateral, -2.40 mm dorsoventral relative to bregma (Josset et al., 2018; Caggiano et al., 2018, van der Zouwen et al. 2021). The ferrule was secured on the cranium using two 00-96×1/16 mounting screws (HRS Scientific, QC, Canada) and dental cement (A-M Systems, Sequim, WA, USA). The scalp incision was sutured, and antibiotics were applied locally.

### *In vivo* optogenetic stimulation

The procedure was as previously reported (van der Zouwen et al. 2021). Briefly, the implanted optic fiber was connected to a 470 nm laser (Ikecool, Anaheim, CA, USA) using a pigtail rotary joint (Thorlabs). The laser was driven with a Grass S88X to generate the stimulation trains (10 s trains, 10 ms pulses, 20 Hz, Caggiano et al. 2018, Josset et al. 2018, van der Zouwen et al. 2021). To visualize optogenetic stimulation, a copy of the stimulation trains was sent to a small (diameter 0.5 cm) low-power (0.13 W) red LED coupled with a 120 MΩ resistance (van der Zouwen et al. 2021). The LED was placed in the field of view of the camera placed above the open field. The 470 nm laser was adjusted to 3.2-30.0% of laser power. The corresponding power measured at the fiber tip with a power meter (PM100USB, Thorlabs) was 0.1–41.2 mW. We estimated irradiance as a function of distance from the optic fiber using a model provided by the Deisseroth lab based on measurements in brain tissues (https://web.stanford.edu/group/dlab/cgi-bin/graph/chart.php, see also Yizhar et al. 2011, Josset et al. 2018). We previously provided evidence that the evoked locomotor responses are specific to blue light in Vglut2-ChR2-EYFP mice, since replacing the blue laser by a red one (589 nm) does not evoke any locomotor response (van der Zouwen et al. 2021).

### Open-field locomotion

The procedure was as previously reported (van der Zouwen et al. 2021). Briefly, locomotor activity was filmed from above in a 40 × 40 cm open field arena at 30 fps using a Canon Vixia HF R800 camera. To measure spontaneous locomotor activity before and after intracerebral injection of the vehicle or 6-OHDA, locomotor activity was recorded during trials of 4.5 min. To measure the effects of optogenetic stimulation, locomotor activity was recorded during trials of 15 min during which 10 stimulation trains were delivered every 80 s at various laser powers. Video recordings were analyzed with DeepLabCut to track user-defined body parts (Mathis et al. 2018, Nath et al. 2019, Hausmann et al. 2021) and a custom Matlab script (Mathworks, Natick, MA, USA) (van der Zouwen et al. 2021). We tracked the body center positions, the corners of the arena for distance calibration, and the small LED to detect optogenetic stimulations. Timestamps were extracted using Video Frame Time Stamps (Matlab File Exchange). Body center positions and timestamps were used to calculate locomotor speed. Body center positions were excluded if their detection likelihood by DeepLabCut was <0.8, if they were outside of the open-field arena, or if body center speed exceeded the maximum locomotor speed recorded in mice (334 cm/s, Garland et al. 1995).

### Limb kinematics

The procedure was as previously reported (van der Zouwen et al. 2021). Briefly, to label hindlimb joints, mice were anesthetized, the hindlimb was shaved and ∼2 mm white dots were drawn on the iliac crest, hip, knee, ankle, and metatarsophalangeal (**MTP**) joints, and toe tip using a fine-tip, oil-based paint marker (Sharpie). After 20 min of recovery from anesthesia, mice were placed in a 1 m long, 8 cm wide transparent corridor. Hindlimb kinematics were recorded at 300 fps from the side using a high-speed Genie Nano Camera M800 camera (Teledyne DALSA, Waterloo, ON, Canada) coupled to a computer equipped with Norpix Streampix software (1st Vision, Andover, MA, USA). For distance calibration, 4 markers (diameter 0.5 cm) were distributed 5 cm apart in the field of view of the camera. To detect optogenetic stimulation, a LED that received a copy of the stimulation trains was placed in the field of view of the camera. Animals were recorded during optogenetic-evoked locomotion and during spontaneous locomotion evoked by a gentle touch of the animal’s tail or a gentle air puff generated by a small air bulb.

The positions of the joints and toe tip were detected using DeepLabCut. A moving average of the MTP speed was used to determine the stance and swing phases by detecting the touchdown and lift-off times with a speed threshold of 9 cm/s, and a minimum of 14 frames above threshold for the lift-off detection (van der Zouwen et al. 2021). The joint positions were used to extract the angles of the hip, knee and ankle joints. The angular variations as a function of time were normalized to step cycle duration using MTP touchdown times as a reference (Leblond et al. 2003).

Frames were excluded from the analysis if the MTPs or any limb joints or the toe tip had a detection likelihood by DeepLabCut was < 0.8, if any paw’s or joint’s speed exceeded 400 cm/s (i.e. maximum locomotor speed of a mouse with a 20% margin to account for increased speed of individual body parts), or if the distance between two adjacent joints was > 2.3 cm (i.e. length of the tibia in wildtype mice) (van der Zouwen et al. 2021).

### DeepLabCut networks

The networks used were the same as those described in van der Zouwen et al. (2021). Briefly, for the analysis of locomotion in the open-field arena, we labelled the body center, the corners of the arena, and the LED to visualize optogenetic stimulation. We used a ResNet-50-based neural network (He et al. 2016, Insafutdinov et al. 2016) with default parameters for 1,030,000 training iterations. We validated with one shuffle and found that the test error was 2.28 pixels and the train error 1.85 pixels (van der Zouwen et al. 2021).

For limb kinematics analysis, we labelled the 5 joints and the toe tip, four distance calibration markers, and the low-power LED to visualize optogenetic stimulation. We used a ResNet-50-based neural network (He et al. 2016, Insafutdinov et al. 2016) with default parameters for 1,030,000 training iterations and one refinement of 1,030,000 iterations. We validated with one shuffle and found that the test error was 2.03 pixels and the train error 1.87 pixels (van der Zouwen et al. 2021).

### Histology

Procedures were as previously reported (Fougère et al. 2021, van der Zouwen et al. 2021). Briefly, mice were anaesthetized using isoflurane (5%, 2.5 L per minute) and transcardially perfused with 30-50 mL of a phosphate buffer solution (0.1M) containing 0.9% of NaCl (PBS, pH = 7.4), followed by 50 mL of PBS solution containing 4% (wt/vol) of paraformaldehyde (PFA 4%). Post-fixation of the brains was performed for 24 h in a solution of PFA 4%. Brains were incubated in a PB solution containing 20% (wt/vol) sucrose for 24 h before histology. Brains were snap frozen in methylbutane (−45°C ± 5°C) and sectioned at -20°C in 40 µm-thick coronal slices using a cryostat (Leica CM 1860 UV). Floating sections at the level of the MLR, striatum and SNc were collected under a Stemi 305 stereomicroscope (Zeiss) and identified using the mouse brain atlas of Franklin and Paxinos (2008).

### Immunofluorescence

The procedure was as previously reported (Fougère et al. 2021, van der Zouwen et al. 2021). All steps were carried out at room temperature unless stated otherwise. The sections were rinsed three times during 10 min in PBS and incubated during 1h in a blocking solution containing 5% (vol/vol) of normal donkey serum and 0.3% Triton X-100 in PBS. The sections were incubated during 48 h at 4°C in a blocking solution containing the primary antibody against TH (rabbit anti-TH, Millipore AB152, lot 2795024 and 3114503 (1:1500) RRID: AB_390204) and gently agitated with an orbital shaker. Then, the sections were washed three times in PBS and incubated during 4 h in a blocking solution containing a secondary antibody to reveal TH (donkey anti-rabbit Alexa Fluor 594, Invitrogen A21207 lot 1890862 and 2145022 (1:400), RRID: AB_141637). The slices were rinsed three times in PBS for 10 min and mounted on Colorfrost Plus (Fisher 1255017) with a medium with DAPI (Vectashield H-1200) or without DAPI (Vectashield H-1000), covered with a 1.5 type glass coverslip and stored at 4°C before observation.

### RNAscope

To detect the *ZsGreen* and *Vglut2* (also called Slc17a6) mRNAs in coronal brain slices, we used the Advanced Cell Diagnostics RNAscope Multiplex Fluorescent Reagent Kit v2 Assay on fixed-frozen tissue samples (ACD 323100-USM). All steps were carried out at room temperature unless stated otherwise. Mice were anesthetized with isoflurane (5%, 2.5 L per minute) and transcardially perfused with 50 mL of a PBS, followed by 50 mL of PBS solution containing 4% (wt/vol) of PFA. Brains were extracted and snap-frozen on dry ice and sectioned at -20°C in 15 µm-thick coronal slices using a cryostat (Leica CM 1860 UV) and mounted onto Colorfrost Plus glass slides (Fisher 1255017). Sections were air-dried during 2 h at -20°C, washed in PBS during 5 min, and baked during 30 min at 60°C in an HybEZ II oven (ACD 321721). Sections were then dehydrated in increasing concentrations of ethanol (50%, 70%, 100%, 100%, 5 min each), treated with hydrogen peroxide during 10 min (ACD 322381, lot 2011534), with RNAscope 1X Target Retrieval Agent during 5 min (ACS 322000, lot 2011356), and with protease III during 30 min in the oven at 40°C (ACD 322381, lot 2011534). Sections were then treated during 2 h at 40°C in the oven with an RNA hybridization antisense probe against ZsGreen-C1 (ACD, 461251, lot 20296A) and another against mouse Slc17a6-C4 (i.e. Vglut2, ACD 319171-C4, lot 21049B). After overnight incubation in a saline sodium citrate solution (175.3g of NaCl and 88.2g of sodium citrate in 1L of distilled water, pH=7.0). C1 probe amplification was done using RNAscope Multiplex Fluorescent Detection Reagents v2 (ACD 323110, lot 2011351). C4 probe amplification was done with the RNAscope Multiplex Fluorescent Detection Reagents v2, with the addition of the HRP-C4 (ACD 323121, lot 2011711). C1 and C4 probes were revealed using Opal Dye 520 (Akoya Biosciences FP1487001KT, lot 201008031, 1:150 or 1:1500) and Opal Dye 690 (Akoya Biosciences FP1497001KT lot 201008030, 1:150 or 1:1500). Sections were counterstained with DAPI (ACD 323108, lot 2011350), mounted with a ProLong Gold Antifade Mountant (Invitrogen P36930, lot 2305164), covered with a 1.5 type glass coverslip and stored at +4°C before observation.

### Microscopy

Brain sections were observed using a Zeiss AxioImager M2 microscope bundled with StereoInvestigator 2018 software (v1.1, MBF Bioscience). To show the expression of ChR2-EYFP in the CnF, high magnification (63X) photographs were taken using a Leica TCS SP8 nanoscope bundled with LASX software (Leica). Composite images were assembled using StereoInvestigator. The levels were uniformly adjusted in Photoshop CS6 (Adobe) to make all fluorophores visible and avoid pixel saturation, and digital images were merged.

### Cell counting

The procedure was as previously reported (Fougère et al. 2021). To estimate the number of TH-positive cells in the SNc, for each animal two coronal brain slices were photographed at 20X magnification with the epifluorescent microscope Zeiss AxioImager M2. On each side of the slice, a region of interest was traced over the photograph using ImageJ, its area was measured and the cells positive for TH were counted and expressed as the number of cells per surface unit (cm^2^). Our criterion for cell count was a clear labeling of the cell body as previously (Fougère et al. 2021).

### TH optical density

To estimate the density of the TH innervation in the striatum, measures of optical density of the TH-immunopositive signal was used (see e.g. Xavier et al. 2005, Perlbarg et al. 2018). For each animal, two coronal sections at the level of the striatum were imaged using the epifluorescent microscope Zeiss AxioImager M2. Using ImageJ, regions of interest were drawn manually over the photographs to delineate the striatum. To estimate optical density in the regions of interest, the photographs were converted into a greyscale and compared to a calibrated greyscale taken from the ImageJ optical density calibration protocol (https://imagej.nih.gov/ij/docs/examples/calibration/) (see also Xavier et al. 2005).

### Statistical analysis

Data are presented as mean ± standard error of the mean (SEM) unless stated otherwise. No statistical method was used to pre-determine sample sizes, which are similar to those used in the field (e.g. Caggiano et al. 2018, van der Zouwen et al. 2021). No randomization or blinding procedure was used. Statistical analysis was done using Sigma Plot 12.0. Parametric analyses were used when assumptions for normality and equal variance were respected, otherwise non-parametric analyses were used. Normality was assessed using the Shapiro-Wilk test. Equal variance was assessed using the Levene test. To compare the means between two dependent groups, a parametric two-tailed paired t-test or a non-parametric Wilcoxon test were used. To compare the means between two independent groups, a two-tailed t-test or a non-parametric Mann-Whitney rank sum test were used. For more than two dependent groups, a parametric one-way analysis of variance (ANOVA) for repeated measures or a non-parametric Friedman ANOVA for repeated measures on ranks were used. ANOVAs were followed by a Student Newman-Keuls post hoc test for multiple comparisons between groups. Linear and sigmoidal regressions between variables, their significance, and the 95% confidence intervals were calculated using Sigma Plot 12.0. Statistical differences were assumed to be significant when *P* < 0.05.

### Data availability

All relevant data are available from the authors.

## RESULTS

### Striatal 6-OHDA damages the striatal DA innervation and SNc DA cells

Vglut2-ChR2-EYFP mice were injected bilaterally in the striatum with 6-OHDA to damage the nigrostriatal pathway and induce deficits in locomotor activity similar to those reported in PD (0.2% ascorbic acid, 0.9% NaCl saline solution, 6-OHDA 5 mg/mL, 1 µL/side). Another group of animals was injected in the striatum with a vehicle (0.2% ascorbic acid, 0.9% NaCl saline solution, 1 µL/side) (see methods) (Fig. 1A). Striatal 6-OHDA injections significantly decreased the presence of fibers positive for tyrosine hydroxylase (**TH**, the rate limiting enzyme for dopamine synthesis) in the striatum, compared to mice injected with vehicle (Fig. 1A-E). Statistical analysis of the optical density of TH labelling in the striatum indicated a -29% decrease in animals injected with 6-OHDA compared to animals injected with vehicle (*P* < 0.001, t-test, 81.0 ± 2.2 in n = 6 vehicle mice vs. 57.8 ± 3.5 in n = 5 6-OHDA mice) (Fig. 1K). In the SNc, striatal 6-OHDA injections decreased the number of TH^+^ cells/cm^2^ by -27% when compared to mice injected with vehicle (*P* < 0.05, t-test, 8995 ± 656 TH^+^ cells/cm^2^ in n = 6 vehicle mice vs. 6561 ± 531 TH^+^ cells/cm^2^ in n = 5 6-OHDA mice) (Fig. 1F-J, L). When pooling the results from animals injected with vehicle or 6-OHDA, we found a positive linear relationship between the number of TH^+^ cells/cm^2^ in the SNc and the striatal TH optical density (R = 0.70, *P* < 0.05, Fig. 1M). In a separate analysis, we compared the results obtained in mice injected with vehicle to those obtained in another group of mice without surgery (intact mice), and we found no difference in the striatal TH optical density (*P* > 0.05, t-test, 81.0 ± 2.2 in n = 6 vehicle mice vs. 86.0 ± 1.6 in n = 5 intact mice) and no difference in the number of TH^+^ cells/cm^2^ in the SNc (*P* > 0.05 t-test, 8995 ± 656 TH^+^ cells/cm^2^ in n = 6 vehicle mice vs. 9789 ± 483 TH^+^ cells/cm^2^ in n = 6 intact mice). Altogether this indicated that striatal injection of 6-OHDA lesioned the nigrostriatal pathway, while striatal vehicle injection had no detectable effect on the anatomical integrity of the nigrostriatal pathway.

**Figure 1.**
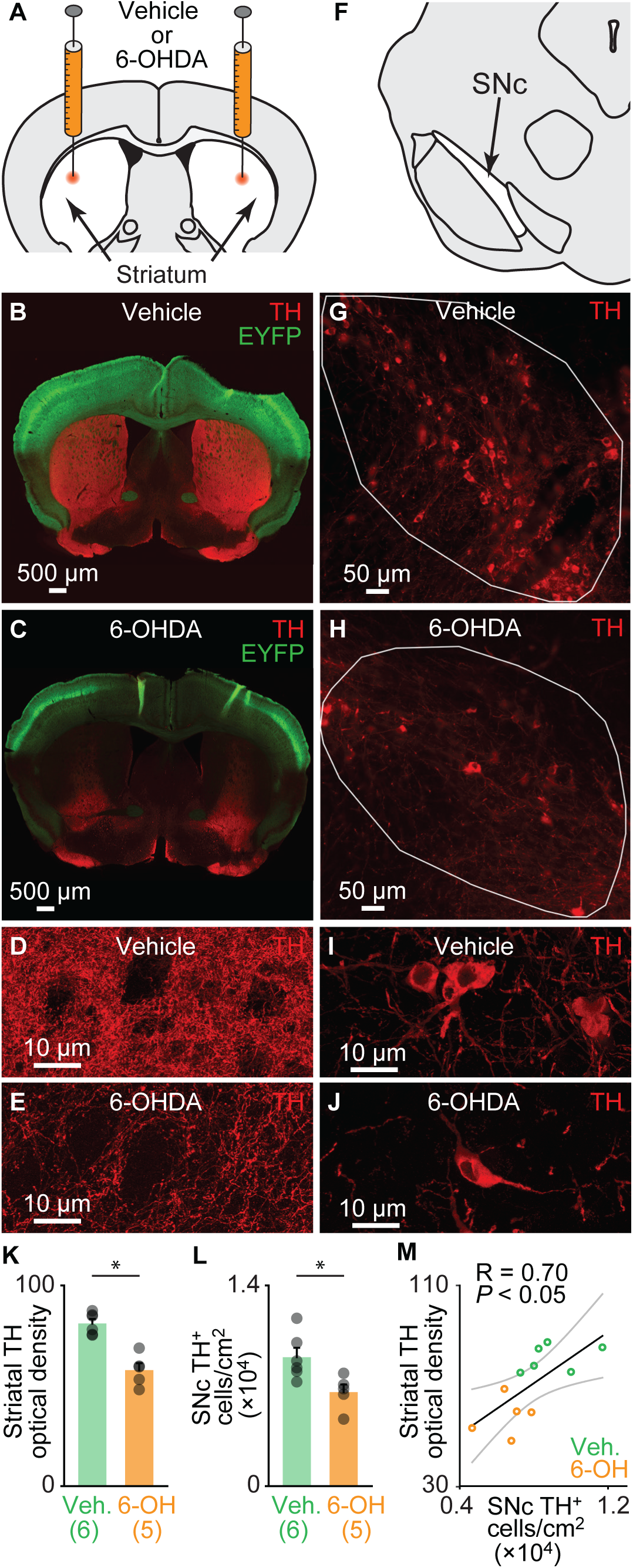
Bilateral striatal injection of 6-hydroxydopamine (6-OHDA) disrupts the nigrostriatal pathway. **A.** Schematic illustration of the injection of 6-OHDA in the striatum (5 mg/mL, 1 µL per side, see methods). **B-C.** Comparison of striatal innervation by tyrosine-hydroxylase (**TH**)-positive fibers in a Vglut2-ChR2-EYFP mouse three days after bilateral striatal injections of vehicle (see methods) (B) and in another one three days after bilateral striatal 6-OHDA injections (C). **D-E.** Magnification showing TH immunofluorescence in the striatum of an animal injected with vehicle (D) or with 6-OHDA (E). **F.** Schematic illustration of the location of TH-positive neurons in the *substantia nigra pars compacta* (**SNc**). **G-H.** Comparison of the presence of the tyrosine-hydroxylase (TH)-positive cells in the SNc (delineated by the solid white lines), in a mouse injected in the striatum with vehicle (G) and another injected with 6-OHDA (H). **I-J.** Magnification showing TH immunofluorescence in the SNc of an animal injected with vehicle (I) or with 6-OHDA (J). **K.** Bar chart illustrating the optical density of TH immunofluorescence in the striatum of mice injected with vehicle vs. mice injected with 6-OHDA. The number of animals used is indicated between brackets. \**P* < 0.05, t-test. **L.** Bar chart illustrating the average bilateral number of TH-positive cells per surface unit in the SNc (two slices counted per mouse, see methods). \**P* < 0.05, t-test. **M.** Relationship between the number of TH-positive cells per surface unit and the striatal TH optical density in the pooled dataset (n = 11 including 5 mice injected with vehicle, green circles, and 6 injected with 6-OHDA, orange circles). \**P* < 0.05, linear fit. The coefficient of correlation (R), its significance (P) and the confidence intervals (grey) are illustrated.

### Striatal 6-OHDA evokes locomotor deficits in the open-field arena

We tracked locomotor activity from above in the open-field arena using video recordings coupled with deep learning-based analysis (see methods) (Fig. 2A-D). When comparing locomotor activity during 4.5 min trials before and after bilateral striatal injection of 6-OHDA (n = 5 mice, 7 trials per animal per condition), we found a decrease of -91% of time spent in locomotion (*P* < 0.01, paired t-test, 64.5 ± 6.5 before vs. 5.9 ± 3.2 s after, Fig. 2E), a decrease of -89% in the number of locomotor initiations (*P* < 0.01, paired t-test, 60.7 ± 6.4 vs. 6.4 ± 3.2 locomotor initiations, Fig. 2F), a decrease of -83% in locomotor speed (*P* < 0.01, paired t-test, 5.1 ± 0.5 vs. 0.9 ± 0.2 cm/s, Fig. 2G), a decrease of -22% in the duration of the locomotor bouts (*P* < 0.05, paired t-test, 1.1 ± 0.0 vs. 0.8 ± 0.1 s, Fig. 2H), and a +29% increase of time spent immobile (*P* < 0.01, paired t-test, 205.4 ± 6.5 vs. 264.0 ± 6.5 s, Fig. 2I).

**Figure 2.**
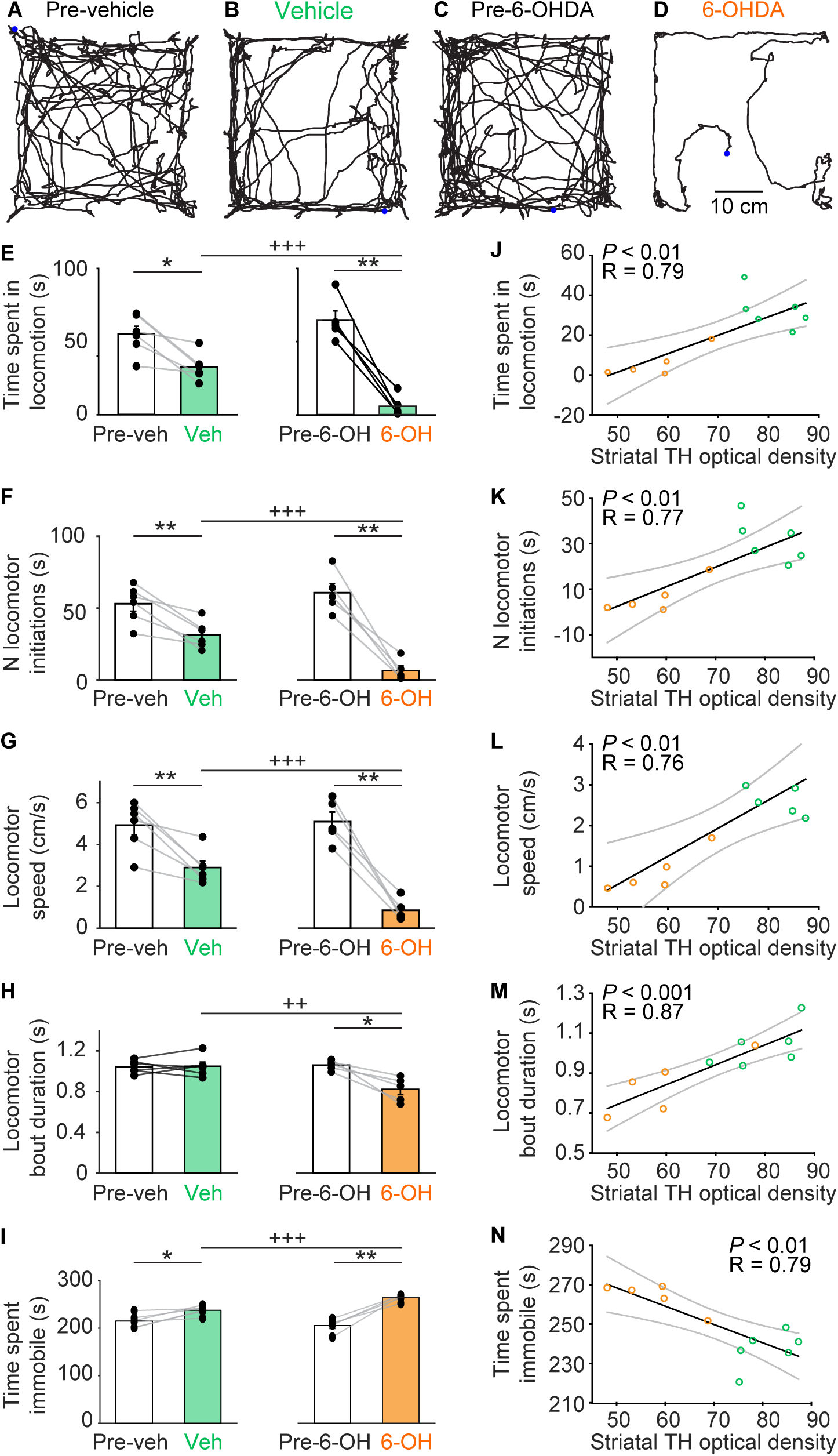
Bilateral striatal injection of 6-hydroxydopamine (6-OHDA) reduces locomotor activity in the open-field arena. **A-D.** Locomotor activity recorded from above in the open-field recorded during a single trial of 4.5 min before and after bilateral striatal injection of vehicle (A-B) or 6-OHDA (C-D). The blue point illustrates the animal’s position at the beginning of the trial. **E-I.** Comparison of locomotor parameters in the open field arena in 6 animals injected in the striatum with vehicle and 5 animals injected with 6-OHDA (7 trials recorded per animal). In E, time spent in locomotion, i.e. time duration during which locomotor speed is higher than 3 cm/s for at least 0.5 s. In F, number (N) of locomotor initiations, i.e. number of times when speed is higher than 3 cm/s for at least 0.5 s. In G, locomotor speed in cm/s. In H, locomotor bout duration, i.e. time duration during which locomotor speed is higher than 3 cm/s for at least 0.5 s. In I, time spent immobile, with immobility defined as total time without locomotion. **J-N.** Linear relationships between the striatal optical density of tyrosine hydroxylase (**TH**) immunofluorescence and the locomotor parameters described in E-I. For each fit, the coefficient of correlation (R), its significance (P) and the confidence intervals (grey) are illustrated. Mice injected with vehicle appear as green circles, those injected with 6-OHDA as orange circles. \**P* < 0.05, \*\**P* < 0.01, \*\*\**P* < 0.001, paired-test or Wilcoxon test; ^++^*P* < 0.01, ^+++^*P* < 0.001, t-test or Mann-Whitney rank sum test.

In comparison, animals injected with vehicle bilaterally in the striatum (n = 6 mice) displayed smaller locomotor deficits in the open-field arena (Fig. 2E-I). Importantly, the locomotor deficits were more dramatic in animals injected with 6-OHDA than in those injected with vehicle. In mice injected with 6-OHDA, we found a shorter time spent in locomotion (*P* < 0.001, t-test, 32.5 ± 3.8 in 6 vehicle mice vs. 5.9 ± 3.2 s in 5 6-OHDA mice, Fig. 2E), a smaller number of locomotor initiations (*P* < 0.001, t-test, 31.5 ± 3.8 vs. 6.4 ± 3.2 locomotor initiations, Fig. 2F), a slower locomotor speed (*P* < 0.001, t-test, 2.9 ± 0.3 vs 0.9 ± 0.2 cm/s, Fig. 2G), a shorter locomotor bout duration (*P* < 0.01, t-test, 1.0 ± 0.0 vs. 0.8 ± 0.0 s, Fig. 2H), and a longer time spent immobile (*P* < 0.001, t-test, 237.5 ± 3.8 vs. 264.0 ± 3.2 s, Fig. 2I). All these parameters were similar in the two groups of mice before injection of 6-OHDA (n = 5 mice) or vehicle (n = 6 mice) (*P* > 0.05 in all cases, t-tests, Fig. 2E-I).

Next, we examined the relationships between the anatomical integrity of the nigrostriatal pathway and the locomotor deficits in the open-field arena by pooling the results of 11 mice injected in the striatum either with vehicle (n = 6 mice) or with 6-OHDA (n = 5 mice). We found positive linear relationships between the striatal TH optical density and: i) the time spent in locomotion (*P* < 0.01, R = 0.79, Fig. 2J), ii) the number of locomotor initiations (*P* < 0.01, R = 0.77, Fig. 2K), iii) the locomotor speed (*P* < 0.01, R = 0.76, Fig. 2L), and iv) the locomotor bout duration (*P* < 0.001, R = 0.87, Fig. 2M). We found a negative linear relationship between the striatal TH optical density and the time spent immobile (*P* < 0.01, R = 0.79, Fig. 2N).

Altogether, these data indicated that striatal 6-OHDA induced major deficits in locomotor activity in the open-field arena. The small locomotor deficits observed in the animals injected with vehicle are likely due to the surgery since vehicle injection did not disrupt the anatomical integrity of the nigrostriatal pathway (see anatomical section above).

### CnF photostimulation evokes locomotion in Vglut2-ChR2-EYFP mice lesioned with 6-OHDA

To determine whether activation of the CnF glutamatergic neurons can increase locomotor activity in PD conditions, we used Cre-dependent expression of transgenes in Vglut2-Cre mice (Vong et al. 2011). To determine whether CnF neurons expressed Vglut2 in our mice, we crossed Vglut2-Cre with ZsGreen-lox mice and performed RNAscope experiments in the offspring. We found that 98.7 % of CnF cells expressing *ZsGreen* mRNA also expressed *Vglut2* mRNA (371/376 cells pooled from n = 4 mice, Fig. 3A-G).

**Figure 3.**
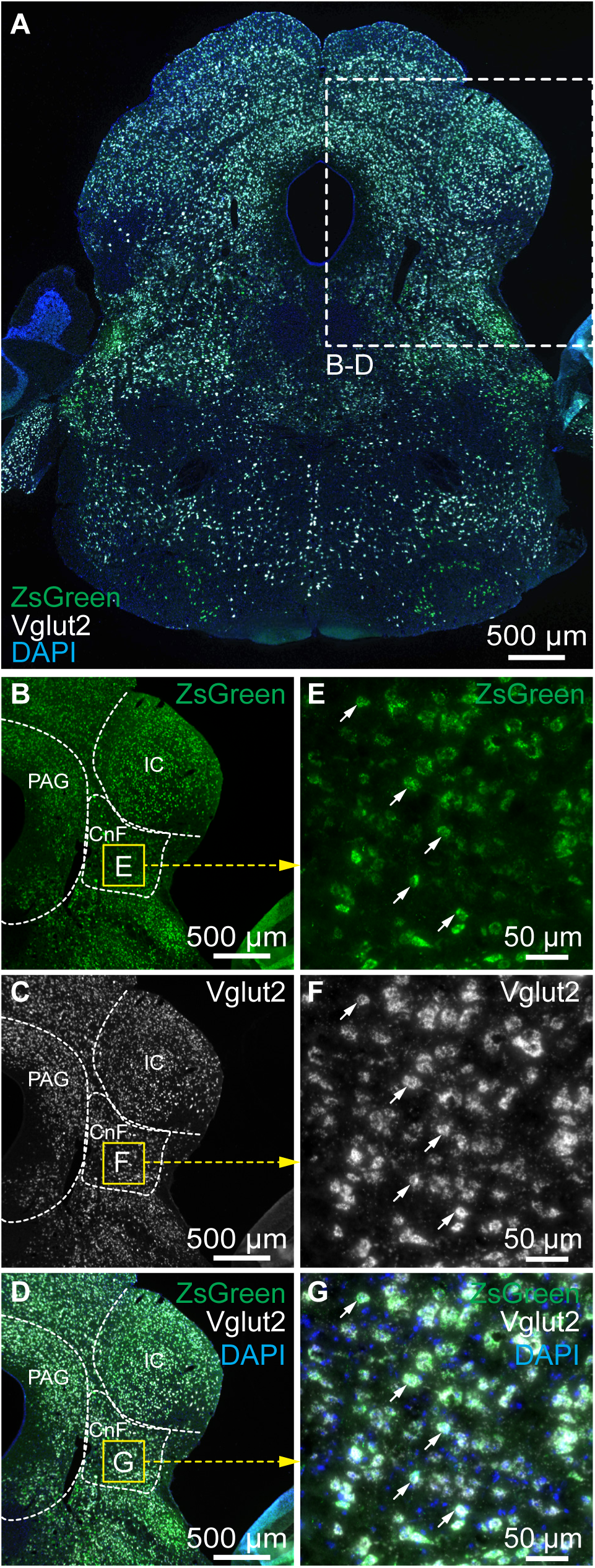
ZsGreen-positive cells in the cuneiform nucleus express the *vesicular glutamatergic transporter 2 (Vglut2)* mRNA in Vglut2-ZsGreen mice. **A.** Coronal brainstem slices from a Vglut2-ZsGreen mouse at the level of the cuneiform nucleus, showing the mRNAs of *ZsGreen* (green), *Vglut2* (white) revealed by RNAscope experiments, and a coloration of the nuclear marker DAPI (blue). **B-D.** Magnification of the slice in A showing the location of the two markers in the cuneiform nucleus. **E-G.** Magnification of the yellow squares in B-D, showing many examples of cells co-expressing *Vglut2* and *ZsGreen* mRNAs (white arrows) in the cuneiform nucleus. CnF, cuneiform nucleus, IC, inferior colliculus, PAG, periaqueductal gray.

Next, to activate CnF glutamatergic neurons with optogenetics, we crossed Vglut2-Cre mice with ChR2-EYFP-lox mice to obtain Vglut2-ChR2-EYFP mice (van der Zouwen et al. 2021). The optic fiber was implanted 500 µm above the right CnF as verified by histology (n = 6 mice; Fig. 4A-B, D), and we validated that ChR2-EYFP was expressed in the CnF (Fig. 4C). Using patch-clamp recordings in brain slices, we previously demonstrated that blue light evoked spikes in MLR neurons at short latency in Vglut2-ChR2-EYFP mice (van der Zouwen et al. 2021). In animals injected bilaterally in the striatum with 6-OHDA, the nigrostriatal pathway was lesioned (Fig. 1) and significant deficits in locomotor activity were observed (Fig. 2). Optogenetic stimulation of CnF Vglut2 neurons with the 470 nm laser evoked robust increases in locomotor activity in the open-field arena in single animal data (Fig. 4F,H) as well as in the pooled data (n = 5 mice, Fig. 4J). Statistical analysis indicated that CnF photostimulation increased the time spent in locomotion during stimulation (Fig. 4L), the number of locomotor initiations (Fig. 4N), the locomotor speed (Fig. 4P), and reduced the time spent immobile (Fig. 4R). Vglut2-ChR2-EYFP mice injected in the striatum with vehicle also displayed increases in locomotor activity when photostimulated in the CnF with the 470 nm laser (Fig. 4E, G, I, K, M, O, Q). Altogether, these data indicate that optogenetic stimulation of CnF Vglut2^+^ neurons can evoke robust increases in locomotor activity in parkinsonian conditions.

**Figure 4.**
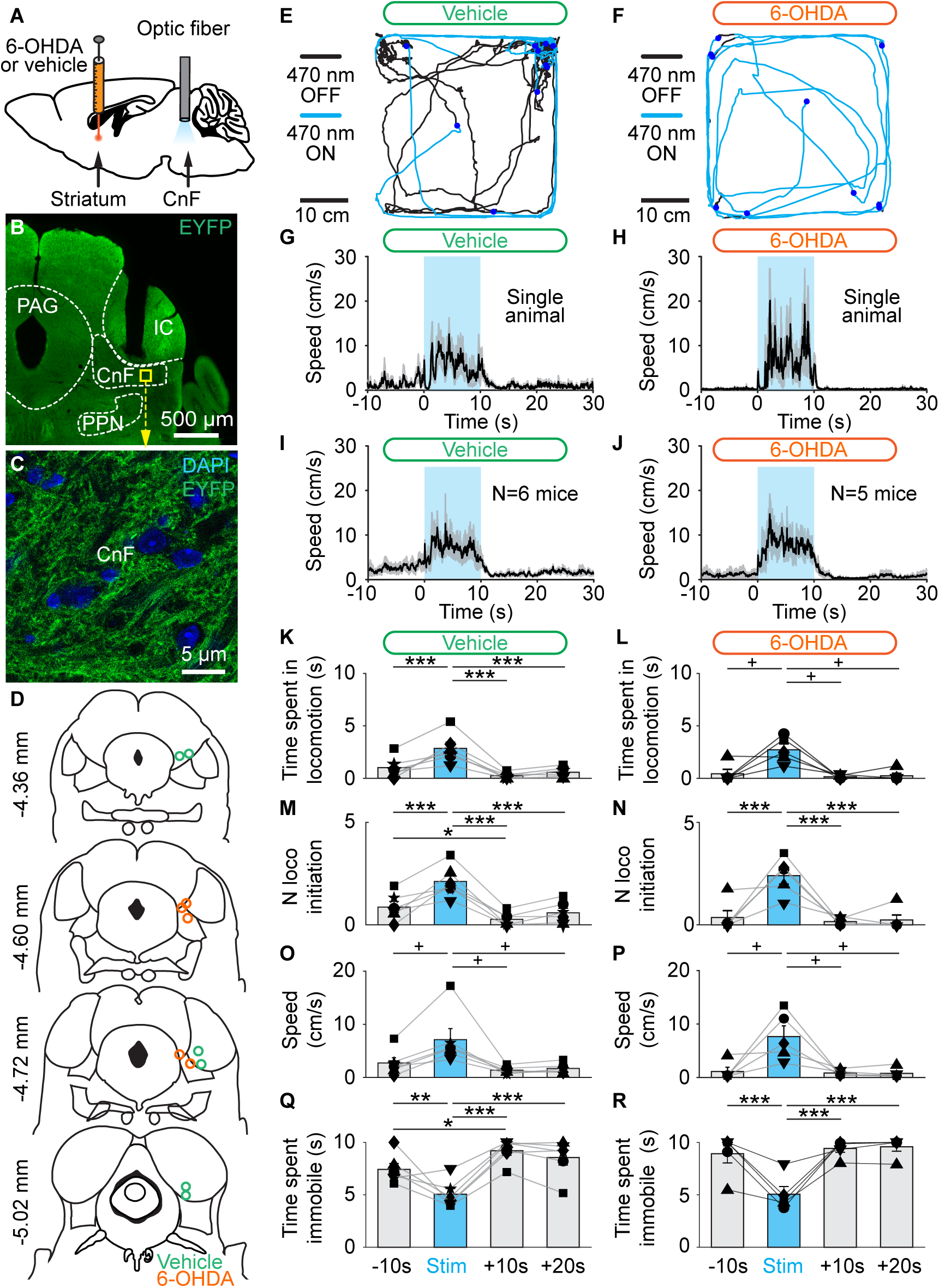
Optogenetic stimulation of the cuneiform nucleus (CnF) increases locomotor activity in Vglut2-ChR2-EYFP mice lesioned with 6-OHDA. **A.** Vglut2-ChR2-EYFP mice were either injected in the striatum with vehicle or 6-OHDA (see methods) and implanted ∼500 µm above the right CnF with an optic fiber. After three days, the effects of optogenetic stimulation of the CnF were tested in the open-field arena. **B.** Photomicrograph showing the position of the optic fiber right above the CnF. **C.** Magnification of the yellow square in B, showing the expression of ChR2-EYFP at the level of the CnF. **D.** Location of the optic fibers after histology of mice injected in the striatum with vehicle (green circles) or 6-OHDA (orange circles). **E-F.** Raw data showing the effects of CnF optogenetic stimulation with a 470 nm laser in a mouse injected in the striatum with vehicle (E, 10 s train, 20 Hz, 10 ms pulses, 5% of laser power), with a 470 nm laser in a mouse injected in the striatum with 6-OHDA (F, 10% of laser power). **G-H.** Locomotor speed (mean ± SEM) before, during and after a 10 s optogenetic stimulation (onset at t=0s) with a 470 nm laser in a single animal (G, 5% of laser power) or in a pool of 6 animals injected in the striatum with vehicle (H, 10 stimulations per animal, 5-20% of laser power). **I-J.** Locomotor speed (mean ± SEM) before, during and after a 10 s optogenetic stimulation (onset at t=0s) with a 470 nm laser in a single animal (I, 10% of laser power) or in a pool of 5 animals injected in the striatum with 6-OHDA (J, 10 stimulations per animal, 5-20% of laser power). **K-R.** Evolution of locomotor parameters before (−10 to 0 s), during (0 to 10 s), and after (10 to 20 s and 20 to 30 s) optogenetic stimulation of the CnF with a 470 nm laser in 6 animals injected in the striatum with vehicle (K, M, O, Q) and in 5 animals injected with 6-OHDA (L, N, P, R). \**P* < 0.05, ** *P* < 0.01, \*\*\**P* < 0.001, Student-Newman-Keuls test after a One-Way ANOVA for Repeated Measures (*P* < 0.001 in K, M, N, Q, R). ^+^*P* < 0.05, Student-Newman-Keuls test after a Friedman Repeated Measures Analysis of Variance on Ranks (*P* < 0.05 in L, P; *P* < 0.01 in O).

### CnF photostimulation controls locomotor speed in Vglut2-ChR2-EYFP mice lesioned with 6-OHDA

A major function of the MLR is to control locomotor speed (for review Ryczko and Dubuc 2013). Next, we determined whether this key property was still functional in PD conditions. In Vglut2-ChR2-EYFP mice injected bilaterally in the striatum with 6-OHDA, increasing the laser power applied during CnF photostimulation increased locomotor speed in single animal data (Fig. 5B). We expressed the laser power and speed as a function of their maximal value per animal and found that speed was controlled by laser power in the data pooled from mice lesioned with 6-OHDA (Fig. 5D). We found a strong sigmoidal relationship between laser power and locomotor speed in mice lesioned with 6-OHDA (R = 0.90, *P* < 0.001, n = 5 mice, Fig. 5F). In Vglut2-ChR2-EYFP mice injected in the striatum with vehicle, we also found that increasing laser power increased locomotor speed in single animal data (Fig. 5A) and in the data pooled from 6 mice (Fig. 5C). We also found in mice injected with vehicle a strong sigmoidal relation between laser power and locomotor speed expressed as a function of their maximal value per animal (R = 0.97, *P* < 0.001, Fig. 5E). To statistically compare the relationships between laser power and speed in mice injected with vehicle or 6-OHDA, we compared the coefficient of correlations of the sigmoidal fits obtained in each animal and found no significant difference (*P* > 0.05, t-test, R = 0.89 ± 0.03 in n = 6 vehicle mice, R = 0.95 ± 0.02 in n = 5 6-OHDA mice). Overall, this suggests that the lesion of the nigrostriatal pathway did not disrupt the ability of CnF Vglut2^+^ neurons to control speed. Small variations in these relationships from one animal to another could be related to the variable depth or antero/posterior level of the implantation sites (Fig. 4D) (van der Zouwen et al. 2021).

**Figure 5.**
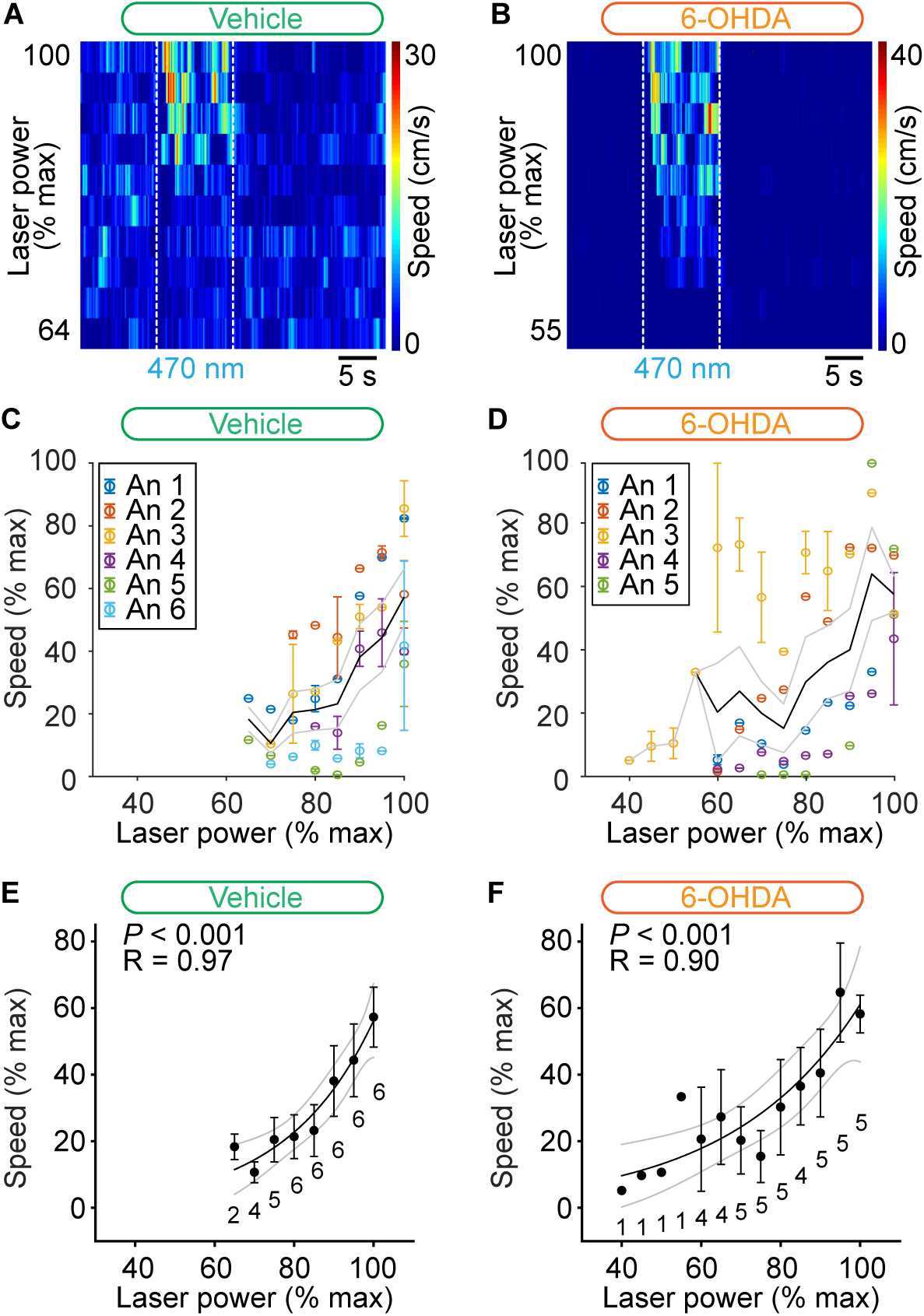
Control of speed by glutamatergic neurons of the cuneiform nucleus (CnF) in Vglut2-ChR2-EYFP mice lesioned with 6-OHDA. **A-B.** Color plots illustrating increases in locomotor speed in the open-field arena during optogenetic stimulation of the CnF in an animal injected in the striatum with vehicle (A, 3.2-5.0% of laser power) or with 6-OHDA (B, 5.5-10.0% of laser power). **C-D.** Locomotor speed (0.3-23.5 cm/s in C, 0.2-30.9 cm/s in D) as a function of laser power (vehicle: 3.2-15.5% of laser power; 6-OHDA: 3.8-20.0% of laser power). Each dot represents the speed (mean ± SEM) measured during 1-3 trials. Speed and laser power were normalized as a function of their maximal value per animal (% max) with a bin size of 5%. The average speed (solid black line) and the SEM (grey solid lines) are illustrated. The data from each mouse is illustrated with a different color. **E-F.** Relationships between locomotor speed (mean ± SEM, bin width 5%) and laser power in the same animals shown in C-D. For each bin, the number of animals is indicated below the data point. The data followed a sigmoidal function both in the 6 animals injected in the striatum with vehicle (E, solid black line, R = 0.97, *P* < 0.001) and in the 5 animals injected with 6-OHDA (F, solid black line, R = 0.90, *P* < 0.001). For each fit, the coefficient of correlation (R), its significance (P) and the confidence intervals (grey) are illustrated.

### Limb kinematics evoked by CnF photostimulation in Vglut2-ChR2-EYFP mice lesioned with 6-OHDA

Next, we determined whether normal locomotion was induced by CnF photostimulation. We recorded the hindlimb kinematics in a linear corridor filmed from the side at high speed (see methods and van der Zouwen et al. 2021). We compared the limb kinematics by measuring the position of each hindlimb joint (iliac crest, hip, knee, ankle, metatarsophalangeal (**MTP**)) and toe over time using DeepLabCut (Fig. 6A-D). In the vehicle group, the angular variations of the hip, ankle and MTP as a function of time were similar during spontaneous locomotion before vehicle injection, and during optogenetic-evoked locomotion after bilateral striatal injection of vehicle (*P* > 0.05 in all cases, paired t-tests, Fig. 6A,B,E,G,I,J), except for the knee joint angle that displayed a slightly higher amplitude (+7%) in optogenetic-evoked locomotion after vehicle injection (49.5 ± 2.3 vs. 53.3 ± 2.7°, *P* < 0.01, paired t-test, n = 4 mice, Fig. 6H). In the 6-OHDA group, the angular variations of the knee, ankle and MTP were similar during spontaneous locomotion before 6-OHDA injection and during optogenetic-evoked locomotion after 6-OHDA injection (P > 0.05 in all cases, paired t-tests, Fig. 6C,D,F,H-J), except for the hip joint angle that displayed a higher amplitude (+38%) in optogenetic-evoked locomotion after 6-OHDA injection (35.1 ± 1.8 vs. 48.7 ± 2.9°, *P* < 0.05, paired t-test, n = 5 mice, Fig. 6G). When comparing vehicle and 6-OHDA groups during spontaneous locomotion prior to intracerebral injections, hip, ankle and MTP angular variations were similar (*P* > 0.05 in all cases, paired t-tests or Mann-Whitney rank sum tests, Fig. 6A,C,E,G,I,J), except for the knee joint angle that showed a higher amplitude (+16%) in the 6-OHDA group (49.5 ± 2.3 vs. 57.7 ± 0.9°, *P* < 0.01, paired t-test, Fig. 6H). Importantly, when comparing limb kinematics during locomotion evoked by optogenetic stimulation in mice injected with vehicle (n = 4 mice) or with 6-OHDA (n = 5 mice), no difference was found in the angular variations of the hip, knee, ankle, or MTP (*P* > 0.05 in all cases, t-tests, Fig. 6G-J). Altogether, this indicated that in parkinsonian conditions, the limb kinematics during locomotion evoked by optogenetic stimulation of CnF Vglut2^+^ neurons were largely similar to those recorded in animals during spontaneous locomotion before lesion.

**Figure 6.**
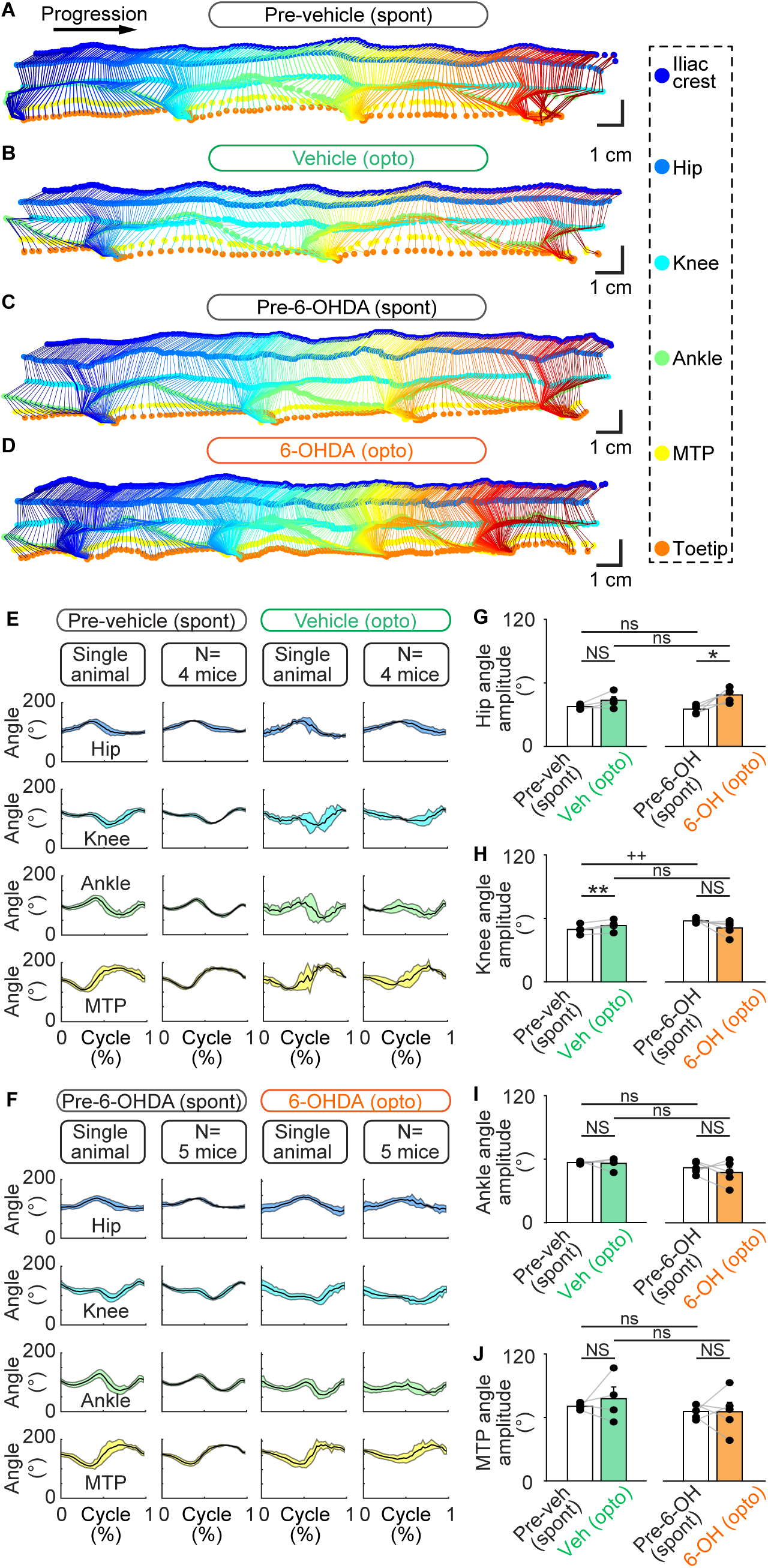
Hindlimb kinematics evoked by optogenetic stimulation of the cuneiform nucleus (CnF) in Vglut2-ChR2-EYFP mice lesioned with 6-OHDA. **A-B.** The movements of six hindlimb joints were tracked from the side at 300 fps in a linear corridor during spontaneous locomotion before striatal injection of vehicle or 6-OHDA, and during optogenetic-evoked locomotion after vehicle injection (C, 6.4% of laser power) or 6-OHDA (D, 30% of laser power). **E-F.** The joint angles at the hip, knee, ankle, and metatarsophalangeal joint (MTP) levels were calculated frame by frame. The cycle was defined as the time duration between two consecutive touchdowns of the MTP. A speed threshold of 9 cm/s was used to define the transitions between swing and stance phases. For single animal data, joint angles (mean ± SD) were plotted for a normalized locomotor cycle during spontaneous locomotion before vehicle (E, 31 steps) or before 6-OHDA injection (F, 30 steps) and during optogenetic-evoked locomotion after striatal vehicle injection (E, 8 steps; 6.4% of laser power) or after striatal 6-OHDA injection (F, 32 steps; 10% of laser power). For the pooled data, joint angles (mean ± SD) were plotted for a normalized locomotor cycle during spontaneous locomotion before vehicle injection (E, 31–39 steps per animal), or before 6-OHDA injection (F, 28–34 steps per animal), during optogenetic-evoked locomotion after vehicle injection (E, 3–19 steps per animal, 5.2–20.0% of laser power) and during optogenetic-evoked locomotion after 6-OHDA injection (F, 3–32 steps per animal, 5–30% of laser power). **F.** Comparison of the amplitude of the hip (G), knee (H), ankle (I), and MTP (J) angles during spontaneous locomotion before vehicle or 6-OHDA injection, and during optogenetic-evoked locomotion after vehicle or 6-OHDA injection (vehicle, n = 6 mice; 6-OHDA, n = 5 mice). NS *P* > 0.05, \**P* < 0.05, \*\**P* < 0.01, paired-test; ns *P* > 0.05, ^++^*P* < 0.01, t-test or Mann-Whitney test.

## DISCUSSION

In the present study, we found that optogenetic stimulation of glutamatergic (Vglut2^+^) neurons in the CnF increases locomotor activity in a 6-OHDA-based mouse model of PD. Bilateral striatal injection of 6-OHDA in Vglut2-ChR2-EYFP mice damaged the TH-positive innervation of the striatum and damaged SNc DA cells. This was associated with major deficits in locomotor activity in the open field arena. Optogenetic activation of Vglut2^+^ neurons in the CnF increased the number of locomotor initiations, reduced the time spent immobile, and locomotor speed was precisely controlled by laser power in mice injected in the striatum with 6-OHDA, as in mice injected with vehicle. Deep learning-based analysis of locomotor movements indicated that limb kinematics evoked by optogenetic stimulation in mice lesioned with 6-OHDA were close to those recorded before lesion. Our study shows that the brainstem locomotor circuits downstream of midbrain DA cells remain functional despite the dramatic decrease in locomotor activity after 6-OHDA lesion. This indicates that CnF Vglut2^+^ neurons are likely a relevant target to improve locomotor function in PD conditions.

### Targeting CnF neurons

Although we cannot completely rule out partial recruitment of Vglut2^+^ neurons of the PPN, or other neighboring structures, our results indicate that the Vglut2^+^ neurons we stimulated are most likely located in the CnF. At the behavioral level, the precise control of speed in freely moving mice that we report here (Fig. 5) is the hallmark of successful activation of glutamatergic neurons in the CnF (Roseberry et al. 2016, Caggiano et al. 2018, Josset et al. 2018, van der Zouwen et al. 2021). The control of locomotor speed exerted by PPN glutamatergic neurons appears to be less robust than that exerted by CnF ones (Caggiano et al. 2018, Josset et al. 2018), making their recruitment unlikely here. We did not record any decelerated locomotor rhythms or locomotor stops that were reported to be induced by activation of glutamatergic PPN neurons in certain cases (Josset et al. 2018, Dautan et al. 2020). At the anatomical level, we carefully verified that the optic fiber tips were located in or above the CnF (Fig. 4D) as in recent studies targeting the same nucleus (Caggiano et al. 2018, Josset et al. 2018, van der Zouwen et al. 2021). The light that could spread down to the PPN would be limited, as we estimated that with the highest laser power value used in the present study, ∼98% of irradiance is lost at 1 mm from the tip of the optic fiber (Yizhar et al. 2011, Josset et al. 2018).

### Targeting glutamatergic neurons

We likely stimulated specifically CnF glutamatergic neurons, as ∼99% of ZsGreen-positive cells in the CnF of Vglut2-ZsGreen mice expressed *Vglut2* mRNA (Fig. 3). This is consistent with the fact that *Vglut2* is specifically expressed in neurons (Li et al. 2013), and with our observation in Vglut2-ZsGreen mice that the vast majority of ZsGreen positive cells are immuno-positive for the neuronal marker NeuN (Fougère et al. 2021, van der Zouwen et al. 2021). Therefore, CnF GABAergic neurons, which decrease locomotor activity when activated (Roseberry et al. 2016, Caggiano et al. 2018), were likely not recruited in our study.

### Clinical relevance

The main message of the present study is that increasing the activity of glutamatergic (Vglut2^+^) neurons in the CnF is a relevant approach to improve locomotor function in PD conditions. Since 2005, the MLR is explored as a DBS target (Plaha and Gill 2005) to improve locomotor function in PD, but the results remain mixed (for review Hamani et al. 2016a,b, Thevathasan et al. 2018). One possible reason is that DBS protocols focused on the PPN but left the CnF largely unexplored, despite the major role of the CnF in locomotor control in mammals (Lee et al. 2014, Roseberry et al. 2016, Caggiano et al. 2018, Capelli et al. 2017, Josset et al. 2018, van der Zouwen et al. 2021, for review Ryczko and Dubuc 2013, Chang et al. 2020). Historically, the MLR was discovered functionally using electrical stimulation in a region encompassing the CnF and the PPN in cats (Shik et al. 1966, for review Ryczko and Dubuc 2013). In humans, careful analysis of DBS electrode position relative to the pontomesencephalic junction revealed that patients in whom DBS was most effective against gait freezing had their electrodes located in or around the CnF (Goetz et al. 2019). Future clinical studies should evaluate the effects of CnF DBS on locomotor function, and a first clinical trial is currently exploring this avenue (NCT04218526, Chang et al. 2021a,c).

Future studies should also aim at understanding how to specifically control glutamatergic (Vglut2^+^) neurons in the CnF in a clinical setting. One possibility would be to consider the electrophysiological properties of CnF glutamatergic neurons to optimize DBS protocols. A recent study indicates that most CnF glutamatergic neurons (86% of Vglut2^+^ cells) have homogeneous properties, whereas PPN glutamatergic neurons are more heterogeneous (Dautan et al. 2020). CnF glutamatergic neurons display membrane potential oscillatory properties in the 20-40 Hz range, whereas PPN glutamatergic neurons in the 10-20 Hz (Dautan et al. 2020). Low electrical stimulation frequencies are needed to gently activate MLR neurons and induce locomotion in animal research [15-50 Hz in cats (Shik et al. 1966, see also Opris et al. 2019), 50 Hz in rats (Bachmann et al. 2013), 20-50 Hz in pigs (Chang et al. 2021b), 5-10 Hz in lampreys and salamanders (Sirota et al. 2000, Brocard et al. 2010, Cabelguen et al. 2003)]. Therefore, clinically, the higher stimulation frequencies (50-140 Hz) used to disrupt the abnormal ongoing rhythmic activity in the subthalamic nucleus should be avoided in the MLR (e.g. Deuschl et al. 2006, Su et al. 2018). A second possibility would be to use pharmacological agents designed to preferentially control the activity of CnF glutamatergic neurons. The team of Garcia-Rill reported that histone deacetylases modulate the electrophysiological activity of PPN neurons in brain slices and PPN oscillations *in vivo* (Urbano et al. 2018, Bisagno et al. 2020). A third possibility would be to develop the use of optogenetics to recruit genetically-defined cell types in humans (Ratner 2021). Two ongoing clinical trials aim at restoring visual detection of in retinis pigmentosa using optogenetics (NCT03326336 and NCT02556736).

An interesting aspect relative to targeting CnF glutamatergic neurons is that their activation should not prevent adaptable navigation through the integration of environmental cues. We recently showed in freely behaving mice that mice can brake and turn during optogenetic stimulation of CnF glutamatergic neurons (van der Zouwen et al. 2021) and this observation holds true in PD animals (Fig. 4E,F). This is consistent with the recent demonstration that separate reticular circuits control speed and turning (Usseglio et al. 2020, Cregg et al. 2020, Lemieux and Bretzner 2019).

### CnF Vglut2^+^ neurons in humans

The presence of glutamatergic and GABAergic neurons in the human CnF has been described using immunohistochemistry (Sebille et al. 2019). The presence of *Vglut2* mRNA has been demonstrated in the human PPN and CnF using *in situ* hybridization (Sebille et al. 2019). It is well established that cholinergic neurons degenerate in the PPN (Hirsch et al. 1987, Jellinger 1988, Karachi et al. 2010, Rinne et al. 2008, Zweig et al. 1987, Karachi et al. 2010). A recent anatomical study reported that some non-cholinergic neurons are also lost in the PPN and CnF in PD, but the status of CnF glutamatergic neurons remains to be studied in detail (Sebille et al. 2019). A key aspect will be to determine whether targeting CnF glutamatergic neurons improves locomotion when the benefits of L-DOPA and subthalamic DBS have worn off (Nonnekes et al. 2015, Gavriliuc et al. 2020). In this regard, it is particularly interesting to underline the preliminary but promising results of a prospective pilot trial performed in Miami, where a patient with severe freezing of gait refractory to L-DOPA therapy showed improvement in locomotor function after bilateral DBS of the CnF (Chang et al. 2021c). This suggests that enough glutamatergic neurons remain responsive in the CnF in parkinsonian conditions to activate downstream locomotor circuits.

## Conclusion

Our work indicates that increasing the activity of CnF glutamatergic (Vglut2^+^) neurons is a relevant approach to improve locomotor function in PD conditions. Future studies should aim at controlling these neurons using pharmacotherapy, optimized DBS protocols, optogenetic or chemogenetic tools, to improve locomotor control and allow smooth navigation in PD conditions.

## ACKNOWLEDGMENTS

We thank Jean Lainé for his technical assistance with the microscopy platform, Florian Bentzinger for providing access to the genotyping equipment. PS is the holder of the Canada Research Chair Tier 1 in the Neurophysiopharmacology of Chronic Pain. This work was supported by the Canadian Institutes of Health Research (407083 to D.R.); the Fonds de la Recherche - Québec (FRQS Junior 1 awards 34920 and 36772 to D.R., Junior 2 award 297238 to D.R.); the Natural Sciences and Engineering Research Council of Canada (RGPIN-2017-05522 and RTI-2019-00628 to D.R.); the Canada Foundation for Innovation (39344 to D.R.); the Centre d’Excellence en Neurosciences de l’Université de Sherbrooke (pilot project grant to D.R.); the Centre de Recherche du Centre Hospitalier Universitaire de Sherbrooke (start-up funding and PAFI grant to D.R.) and the Faculté de médecine et des sciences de la santé (start-up funding to D.R.).

## Notes

**CONFLICT OF INTEREST.** The authors declare no competing interests.

### Competing Interest Statement

The authors have declared no competing interest.

